# Mechanisms governing protective pregnancy-induced adaptions of the pelvic floor muscles in the rat pre-clinical model

**DOI:** 10.1101/2021.08.01.454675

**Authors:** Mary M. Rieger, Michelle Wong, Lindsey A. Burnett, Francesca Boscolo Sesillo, Brittni B. Baynes, Marianna Alperin

**Affiliations:** San Diego, California. Department of Obstetrics, Gynecology, and Reproductive Sciences, University of California, San Diego; San Diego, California. Department of Obstetrics, Gynecology, and Reproductive Sciences, Division of Female Pelvic Medicine and Reconstructive Surgery, University of California, San Diego

**Keywords:** sarcomerogenesis, pelvic floor muscles, birth injury, rat, pregnancy, adaptations

## Abstract

**Background:** The intrinsic properties of pelvic soft tissues in women who do and do not sustain birth injuries are likely divergent, however little is known about this. Rat pelvic floor muscles undergo protective pregnancy-induced structural adaptations, sarcomerogenesis and increase in intramuscular collagen content, that protect against birth injury.

**Objectives:** We aimed to test the following hypotheses: 1) increased mechanical load of gravid uterus drives antepartum adaptations; 2) load-induced changes are sufficient to protect pelvic muscles from birth injury.

**Study Design:** Independent effects of load uncoupled from hormonal milieu of pregnancy were tested in 3- to 4-month-old Sprague-Dawley rats randomly divided into four groups, N=5- 10/group: (1) load^-^/pregnancy hormones^-^ (controls); (2) load^+^/pregnancy hormones^-^; (3) reduced load/pregnancy hormones^+^; (4) load^+^/pregnancy hormones^+^. Mechanical load simulating a gravid uterus was simulated by weighing uterine horns with beads similar to fetal rat size and weight. Reduced load was achieved by unilateral pregnancy after unilateral uterine horn ligation. To assess acute and chronic phases required for sarcomerogenesis, rats were sacrificed at 4 hours or 21 days post bead loading. Coccygeus, iliocaudalis, pubocaudalis and non-pelvic tibialis anterior were harvested for myofiber and sarcomere length measurements. Intramuscular collagen content was assessed using hydroxyproline assay. Additional 20 load^+^/pregnancy hormones^-^ rats underwent vaginal distention to determine whether load-induced changes are sufficient to protect from mechanical muscle injury in response to parturition-associated strains of various magnitude. Data, compared using two-way repeated measures analysis of variance/pairwise comparisons, are presented as mean ± standard error of mean.

**Results:** Acute increase in load resulted in significant pelvic floor muscle stretch, accompanied by acute increase in sarcomere length compared to non-loaded control muscles (coccygeus: 2.69±0.03 vs 2.30±0.06 µm, *P*<0.001; pubocaudalis: 2.71±0.04 vs 2.25±0.03 µm, *P*<0.0001; iliocaudalis: 2.80±0.06 vs 2.35±0.04 µm, *P*<0.0001). After 21 days of sustained load, sarcomeres returned to operational length in all pelvic muscles (*P*>0.05). However, the myofibers remained significantly longer in load^+^/pregnancy hormones^-^ compared to load^-^ /pregnancy hormones^-^ in coccygeus (13.33±0.94 vs 9.97±0.26 mm, *P*<0.0001) and pubocaudalis (21.20±0.52 vs 19.52±0.34 mm, *P*<0.04) and not different from load^+^/pregnancy hormones^+^ (12.82±0.30 and 22.53±0.32mm, respectively, *P*>0.1), indicating that sustained load induced sarcomerogenesis in these muscles. Intramuscular collagen content in load^+^/pregnancy hormones^-^ group was significantly greater relative to controls in coccygeus (6.55±0.85 vs 3.11±0.47µg/mg, *P*<0.001) and pubocaudalis (5.93±0.79 vs 3.46±0.52 µg/mg, *P*<0.05) and not different from load^+^/pregnancy hormones^+^ (7.45±0.65 and 6.05±0.62 µg/mg, respectively, *P*>0.5). Iliocaudalis required both mechanical and endocrine cues for sarcomerogenesis. Tibialis anterior was not affected by mechanical or endocrine alterations. Despite equivalent extent of adaptations, load-induced changes were only partially protective against sarcomere hyperelongation.

**Conclusions:** Load induces plasticity of the intrinsic pelvic floor muscle components that renders protection against mechanical birth injury. The protective effect, which varies between individual muscles and strain magnitudes, is further augmented by the presence of pregnancy hormones. Maximizing impact of mechanical load on pelvic floor muscles during pregnancy, such as with specialized pelvic floor muscle stretching regimens, is a potentially actionable target for augmenting pregnancy-induced adaptations to decrease birth injury in women who may otherwise have incomplete antepartum muscle adaptations.

**AJOG at a Glance:**

A. Why was the study conducted?
  - To determine the role of mechanical load, uncoupled from the hormonal milieu of pregnancy, in driving protective pregnancy-induced adaptations previously discovered in the rat pelvic floor muscles.
B. What are the key findings?
  - Mechanical load, in the absence of pregnancy hormones, induces sarcomerogenesis and extracellular matrix remodeling in rat pelvic floor muscles.
  - Load-induced adaptations are partially protective against mechanical pelvic floor muscle injury consequent to parturition-associated strains.
C. What does this study add to what is already known?
  - The effect of sustained increased mechanical load, uncoupled from the hormonal milieu of pregnancy, on pelvic floor muscle plasticity has not been previously studied.
  - Modulating pelvic floor muscles’ stretch antepartum, such as with specialized pelvic floor physical therapy regimens, could be a promising approach for augmentation of protective muscle adaptations in women.

## INTRODUCTION

Pelvic floor disorders (PFDs), including pelvic organ prolapse, urinary incontinence and fecal incontinence, are highly prevalent conditions that adversely impact the quality of life of women. Dysfunction of the pelvic floor muscles (PFMs), which include the three paired muscles – pubovisceralis and iliococcygeus that comprise levator ani, and coccygeus – and specifically levator ani has been implicated as one of the key risk factors in the pathogenesis of PFDs.^1,2^ Vaginal childbirth is an inciting event for pelvic floor dysfunction in many women, in part because parturition results in elongation of the PFMs up to 300% of their resting muscle length.^3^ These dramatic strains substantially exceed the 60% elongation that has been shown to result in reproducible injury in limb skeletal muscles.^3-5^ Curiously, for reasons yet unknown, a large proportion of vaginally parous women do not develop pelvic floor dysfunction despite similar obstetrical variables to women who do have such dysfunction postpartum.^6^

PFMs are skeletal muscles composed of myofibers that are, in turn, made of muscle basic contractile units - sarcomeres. Skeletal muscles exhibit plasticity in response to alterations in physiological cues,^7^ including the dynamic assembly of the sarcomere units, known as sarcomerogenesis. In the limb, when muscles are subjected to increased mechanical load, the sarcomeres acutely elongate in response to muscle stretch.^8^ If the muscle stretch is sustained, sarcomeres are added in series to restore operational sarcomere length, resulting in increased fiber length and facilitating optimal *in vivo* muscle function.^5,9,10^ Another important structural component of skeletal muscles is the intramuscular connective tissue network that surrounds the contractile myofibers. The intramuscular extracellular matrix (ECM), primarily composed of collagens, provides support to myofibers and bears the majority of muscle’s passive load. Both *in vitro* and *in vivo* studies suggest that mechanical loading induces ECM remodeling in the limb muscles via growth-factor-mediated cell-signaling pathways, presumably to ensure mechanical stability of the muscle fibers; however, this process is not well understood.^11,12^

Significant technical and ethical constraints preclude direct studies of the human PFMs, especially in pregnant women. Consequently, we utilize a validated rat model, as the rat PFM anatomy and architecture are well suited for the studies of the human PFMs.^13,14^ In response to the physiological cues associated with pregnancy, the rat vagina and supportive tissues exhibit significant plasticity which facilitate the ability of the vagina to withstand parturition.^15-17^ We have previously shown that the rat PFMs also undergo adaptations during pregnancy, specifically myofiber elongation via sarcomerogenesis.^18^ This allows PFMs to maintain operational sarcomere length and preserves muscle force generation capacity necessary to support the increased load of the pregnant uterus. Additionally, sarcomerogenesis appears to protect against PFM injury during parturition because of increased ability to withstand muscle strain without pathological sarcomere hyperelongation and the associated myofibrillar disruption.^19^ Pregnancy-induced alterations also take place in PFMs’ ECM. We found that, in contrast to other pelvic tissues, intramuscular collagen content significantly increases in PFMs in pregnancy.^20^ This increase in ECM collagen content is presumed to stabilize elongated PFM fibers and to further protect them from overstretching during parturition. However, the mechanisms leading to sarcomerogenesis and ECM remodeling during pregnancy remain unknown.

In the current study, using the rat pre-clinical model validated for the investigations of human PFMs^13,14^, we aimed to: 1a) determine the acute effect of increased mechanical load on PFMs; 1b) decipher the relative contributions of the chronic mechanical load and pregnancy hormonal milieu on the pregnancy-induced adaptations of PFMs; and 2) assess whether load-induced alterations modulate PFMs’ response to parturition-associated strains. We hypothesized that acute increase in mechanical load imposed on PFMs would result in muscle and sarcomere stretch. We opined that this acute change in sarcomere length (L_s_), combined with sustained increased load, would be sufficient to induce sarcomerogenesis and increased ECM collagen content in PFMs observed during pregnancy. Finally, we hypothesized that load-induced sarcomerogenesis and increased ECM collagen content are sufficient to protect PFMs from the intrapartum sarcomere hyperelongation, the major cause of mechanical muscle injury, in the absence of other physiologic changes of pregnancy.

## MATERIALS AND METHODS

To delve into the potential mechanisms that govern protective PFM adaptations, we started by uncoupling the endocrine and mechanical effects of pregnancy on the muscle structural parameters. To segregate mechanical load from the hormonal cues, we exposed PFMs to the following <u>*four sets of conditions*</u>: 1) load^-^/pregnancy hormones^-^ (non-pregnant control); 2) load^+^/pregnancy hormones^-^ (non-pregnant loaded); 3) reduced load/pregnancy hormones^+^ (unilateral pregnancy); and 4) load^+^/pregnancy hormones^+^ (pregnant) (Figure 1). Group 3 was chosen as a model of reduced load/pregnancy hormones^+^ because it is not possible to recapitulate the full array of complex hormonal alterations that occur in pregnancy without exposing PFMs to at least some increased load, as the hormonal milieu is naturally induced by the conceptuses themselves.

**Figure 1:**
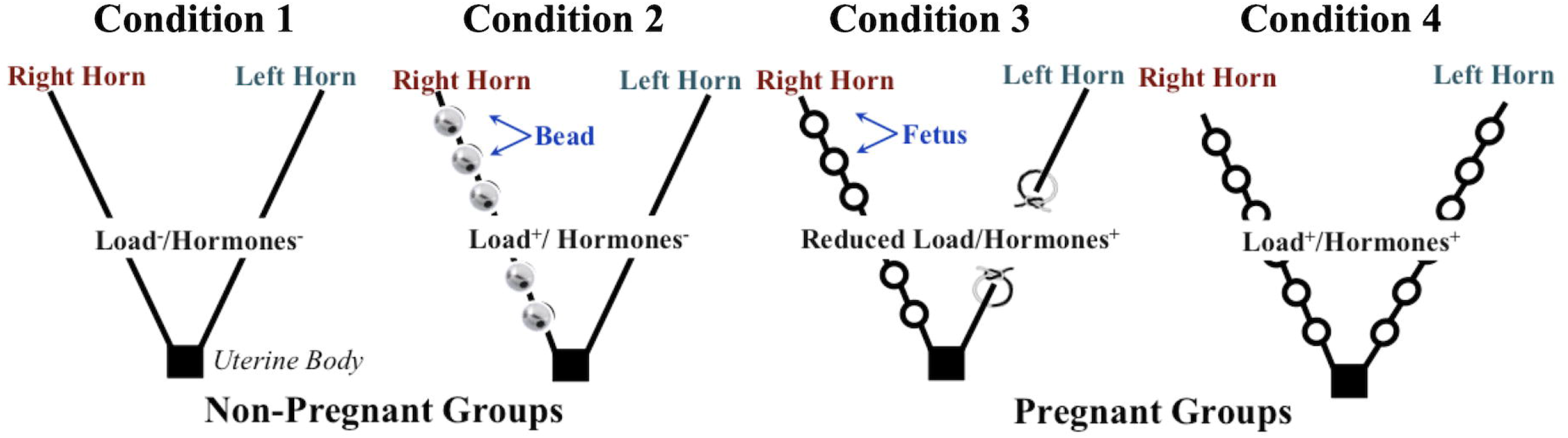
A schematic diagram of experimental approach. To determine the independent and combined roles of mechanical load and hormonal milieu of pregnancy in driving pregnancy-induced adaptations in the rat pelvic floor muscles (PFMs), structural muscle parameters were compared between PFMs subjected to four sets of conditions: 1) load^-^/pregnancy hormones^-^; 2) load^+^/pregnancy hormones^-^; 3) reduced load/pregnancy hormones^-^; and 4) load^+^/pregnancy hormones^+^.

The University of California San Diego Institutional Animal Care and Use Committee (IACUC) approved all study procedures. Three- and 4-month-old Sprague-Dawley rats (Envigo, Indianapolis, IN) were used in the following series of experiments. Rats were housed 2-3/cage according to the IACUC standards and allowed ad lib access to food and water.

### Non-pregnant rat model of mechanical loading of the pelvic floor muscles

To simulate the increased load imposed on PFMs during pregnancy without the concomitant effects of hormonal alterations, we developed a novel non-pregnant rat model with a weight-loaded uterine horns (Figure 1). Three-month-old nulligravid rats (N=10) were anesthetized with isofluorane and administered a pre-operative subcutaneous injection of buprenorphine sustained release at a dose of 1.0 mg/kg. The abdominal fur was removed with depilatory cream (Nair Hair Remover, Ewing, NJ), the skin was sterilized with 4% chlorhexidine gluconate solution (Hibiclens, Norcross, GA), and rats were sterilely draped. Midline laparotomy was performed using standard aseptic techniques. One of the two uterine horns, chosen at random, was exteriorized, and six 3-gm sterile stainless-steel beads (Bullet Weights, Alda, NE), each similar in size and weight to a late pregnant rat fetus, were attached to the anti-mesenteric border using silk sutures (Ethicon, Somerville, NJ). The number of stainless-steel beads was chosen because six fetuses is the median number of conceptuses in each uterine horn during spontaneous rat pregnancy.^17^ The weight-loaded uterine horn was returned into the peritoneal cavity. The fascia was closed with 3-0 polyglactin 910 suture (coated VICRYL, Ethicon, Somerville, NJ) in a continuous running fashion, and the skin was closed with the same suture material in a continuous subcuticular fashion. Animals received an immediate post-operative subcutaneous intra-incisional injection of 0.25% bupivacaine at a dose of 0.4 mL/kg. Animals were euthanized either 4 hours post loading, to assess acute effect of mechanical load on PFMs, or 21 days later, to simulate the sustained load until late gestation.

In our initial perturbation of the model, we hoped to capitalize on the bi-horn rat uterine anatomy, with contralateral PFMs within one animal representing two sets of conditions: load^-^ /pregnancy hormones^-^ (side of non-loaded horn, control) and load^+^/pregnancy hormones^-^ (side with horn loaded with beads, non-pregnant loaded). Interestingly, the fiber length comparisons between contralateral PFMs revealed that sustained load affected both sides, with no significant differences in fiber lengths identified between sides for either of the PFM pairs examined (*P*>0.5, Supplemental Table 1). Thus, we selected rats with unilaterally weighed uterine horns to represent the load^+^/pregnancy hormones^-^ condition, with a separate group of non-pregnant unperturbed rats used to represent load^-^/pregnancy hormones^-^ condition. In addition, fiber length in non-pregnant rats with single loaded uterine horn for 21 days did not differ from that in late pregnant animals. This is likely because beads equal in weight and size to term fetal rats were attached for the entire duration. Nevertheless, to avoid the risk of exceeding adaptations observed in pregnancy, we proceeded with unilateral loading in the load^+^/pregnancy hormones^-^ group.

### Pregnant rat model of reduced loading of the pelvic floor muscles

To create the physiological state of pregnancy with reduced mechanical load, 3-month-old nulligravid rats (N=9) were subjected to a unilateral horn ligation, right or left side selected at random. The selected uterine horn was exteriorized through the laparotomy incision, as described above, and two silk sutures (Ethicon, Somerville, NJ) were placed approximately 1.5 cm apart and tied down. The intervening portion of the horn was excised (Figure 1), and the uterine horn was returned to the peritoneal cavity. The incisions were closed as described above. After 5 days of recovery, the rats were mated and examined daily. The day the vaginal plug was observed was designated as gestational day 1. The animals were euthanized on gestational day 21 (late pregnant).

### The effect of sustained mechanical load on the pelvic floor muscles’ response to parturition-associated strains

To determine whether sustained exposure to increased load in the absence of pregnancy hormonal milieu impacts PFM response to parturition-associated strains, 3-month-old nulligravid rats (N=20) underwent the mechanical loading, as described above. Animals were housed for 21 days. On day 21 after loading, the rats underwent an established vaginal balloon distension procedure.^19^ Two volumes representing physiologic (3 ml, well-approximates fetal rat size) and supraphysiologic (5 ml, approximately 67% larger than fetal rat size) strains were tested (N=10/volume).^19^ Animals were euthanized after 2 hours of vaginal distension.

### Muscle Architectural Parameters

The rat coccygeus and the two components of levator ani (pubocaudalis and iliocaudalis)^21^, as well as tibialis anterior that served as non-pelvic control muscle were fixed *in situ* in formaldehyde for 3-5 days after euthanasia to preserve *in vivo* muscle architecture. Muscle length (L_m_) was measured *in situ* using digital calipers, after which bilateral PFMs and tibialis anterior were harvested and microdissected for fiber length (L_f_) measurement and high-throughput L_s_ measurement by laser diffraction using validated methods.^18,22^

### Intramuscular Extracellular Matrix Assessment

Hydroxyproline, a major component of collagen, was measured to determine the intramuscular ECM content using a validated protocol.^20,23^ Samples were procured from the mid-belly of PFMs and tibialis anterior (3 samples/each muscle), weighed, and hydrolyzed in 6 N hydrochloric acid at 110 ºC for 24 hours. Experimental samples were placed into the 96-well plates in duplicate along with the standards and incubated with a chloramine-T solution, followed by the addition of a p-dimethylaminobenzaldehyde solution. We used spectrophotometry at 550 nm to determine hydroxyproline concentration, normalized to the wet weight of the sample, and converted to collagen using the constant of 7.46, the number of hydroxyproline residues per collagen molecule.

### Statistical Analysis

Structural parameters of each individual PFM subjected to variable conditions, illustrated in Figure 1, and tibialis anterior were compared using 2-way repeated measures ANOVA (factors: load/hormonal status x muscle). Sample size was explored for the key variables of interest (normalized fiber length (L_fn_)^18^, sarcomere length (L_s_)). We set type I error α = 0.05, power (1-β) = 0.80. Based on Cohen’s d effect size of 3.2, power calculation (G*Power) yielded n=4 animals/group. ^24^ We increased the sample size in the animals subjected to surgical procedure (load^+^/hormones^-^ and reduced load/hormones^+^) to account for potential attrition due to post-operative complications. Given a large effect size for collagen content in our previous studies^18^, this sample size was sufficient for this outcome of interest. For L_s_ changes in response to vaginal distention, n=4-5 rats/group/volume was needed for coccygeus and pubocaudalis, given the large effect size, and n=10/group/volume was needed for iliocaudalis that experiences less strain^19^. Post-hoc pairwise comparisons, when appropriate, were conducted with tests adjusted for multiple comparisons. All data were checked for normality to satisfy the assumptions of the parametric tests. All analyses were performed with GraphPad Prism 9.1.1, CA, USA.

## RESULTS

### The acute effect of increased mechanical load on the pelvic floor muscles’ contractile myofibers

The increase in mechanical load induced by uterine bead loading (Figure 2A) resulted in immediate PFM stretch, evidenced by significantly increased muscle length (L_m_) of each PFM on the loaded side compared to the contralateral non-loaded side and unperturbed PFMs in non-pregnant animals (Figure 2B and 2C). In the acute setting, an increase in sarcomere length (L_s_) on the loaded side was similarly observed (Figure 2D). These results demonstrate that increased load imposed on PFMs by the weighted uterine horn leads to the acute whole muscle stretch and the paralleling sarcomere elongation, indicated by increased L_s_. In appendicular muscles, increase in L_s_ serves as a strong impetus for sarcomerogenesis in the face of continued exposure to mechanical load.^3^ Thus, we next proceeded to assess whether load-induced sarcomerogenesis takes place in PFMs in the non-pregnant model subjected to the sustained load, such as that observed in pregnancy.

**Figure 2:**
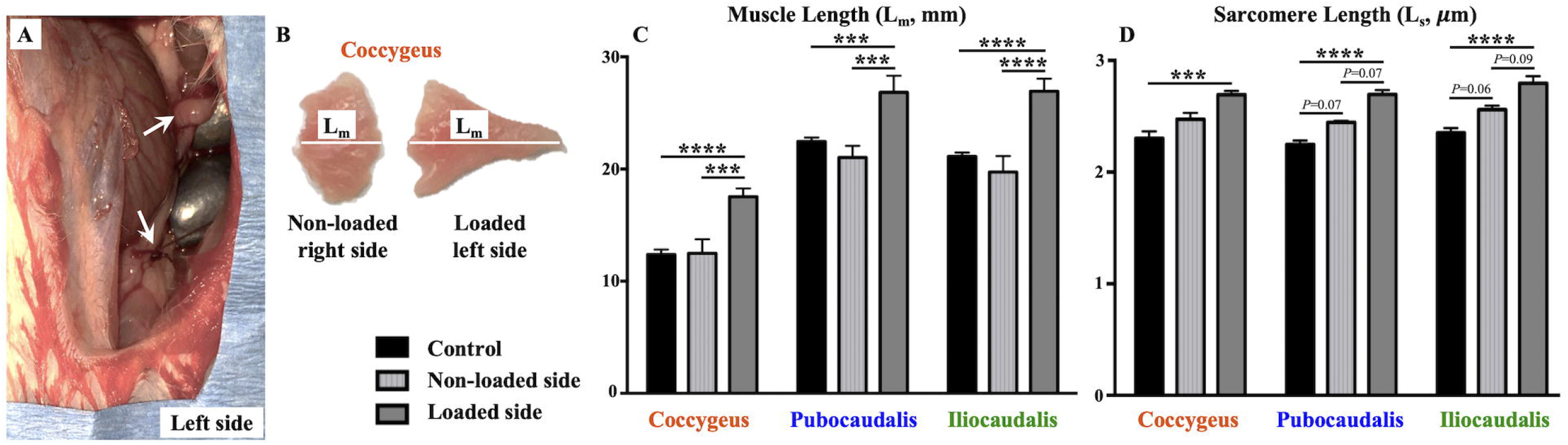
Acute impact of increased mechanical load on the pelvic floor muscles’ (PFMs) contractile myofibers. (**A**) Left uterine horn in non-pregnant rat, loaded with 3-gram stainless steel beads (arrows), each similar in size and weight to a late pregnant rat fetus, visible through a laparotomy incision. (**B**) Bilateral coccygeus muscles, with significantly increased muscle length (L_m_) of the left muscle (loaded side), relative to the right muscle. (**C**) Acute changes in the muscle lengths (in millimeters) of the PFMs subjected to increased load, relative to non-loaded contralateral PFMs and unperturbed controls. N=4/group. (**D**) Acute changes in the sarcomere lengths (in micrometers) of the PFMs subjected to increased load, relative to non-loaded contralateral PFMs and unperturbed controls. N=4/group. *Footnote:* Data are presented as mean ± standard error of mean. \**P* < 0.05; \*\**P* < 0.01; \*\*\**P <* 0.001; \*\*\*\**P* < 0.0001 derived from repeated measures two-way analysis of variance followed by pairwise comparisons with Tukey’s test.

### The effect of sustained mechanical load, uncoupled from the endocrine milieu of pregnancy, on the PFMs’ structural parameters

In skeletal muscles, fiber length (L_f_) can change secondary to either 1) sarcomere elongation/contraction or 2) adaptive assembly/disassembly of the sarcomeres. Thus, we first examined L_s_ of PFMs and tibialis anterior. There was no difference in L_s_ between any of the experimental conditions (load^-^/pregnancy hormones^-^, load^+^/pregnancy hormones^-^, reduced load/pregnancy hormones^-^, and load^+^/pregnancy hormones^+^) for all muscles examined (Table 1). These results mean that any fiber elongation would be the result of adaptive sarcomerogenesis due to sustained load rather than persistent sarcomere stretch observed in the acute phase.

**Table 1.** Sarcomere length (in micrometers) of the pelvic floor muscles and tibialis anterior in the presence of load and/or pregnancy hormones presented as mean ± standard error of mean

| Muscle | Load<sup>-</sup>/<br>Hormones<sup>-</sup><br>(N=10) | Load<sup>+</sup>/<br>Hormones<sup>-</sup><br>(N=9) | Reduced Load/<br>Hormones<sup>-</sup><br>(N=9) |  | Load<sup>+</sup>/<br>Hormones<sup>+</sup><br>(N=5) |  |
| --- | --- | --- | --- | --- | --- | --- |
| Coccygeus | 2.38 ± 0.05 | 2.36 ± 0.05 | 2.38 ± 0.03 |  | 2.26 ± 0.04 |  |
| Pubocaudalis | 2.36 ± 0.03 | 2.32 ± 0.05 | 2.39 ± 0.04 |  | 2.27 ± 0.03 |  |
| Iliocaudalis | 2.46 ± 0.03 | 2.36 ± 0.04 | 2.46 ± 0.03 |  | 2.40 ± 0.05 |  |
| Tibialis Anterior | 2.60 ± 0.02 | 2.60 ± 0.02 | 2.64 ± 0.02 |  | 2.54 ± 0.01 |  |
|  | *P*-value<sup>*</sup> |  |  |  |  |  |
| Muscle | Load<sup>-</sup>/<br>Hormones<sup>-</sup><br>vs<br>Load<sup>+</sup>/<br>Hormones<sup>+</sup> | Load<sup>-</sup>/<br>Hormones<sup>-</sup><br>vs<br>Load<sup>+</sup>/<br>Hormones<sup>-</sup> | Load<sup>-</sup>/<br>Hormones<sup>-</sup><br>vs<br>Reduced<br>Load/<br>Hormones<sup>+</sup> | Load<sup>+</sup>/<br>Hormones<sup>+</sup><br>vs<br>Load<sup>+</sup>/<br>Hormones<sup>-</sup> | Load<sup>+</sup>/<br>Hormones<sup>+</sup><br>vs<br>Reduced<br>Load/<br>Hormones<sup>+</sup> | Load<sup>+</sup>/<br>Hormones<sup>-</sup> vs<br>Reduced<br>Load/<br>Hormones<sup>+</sup> |
| Coccygeus | 0.06 | 0.98 | 0.99 | 0.17 | 0.09 | 0.99 |
| Pubocaudalis | 0.21 | 0.72 | 0.95 | 0.82 | 0.07 | 0.42 |
| Iliocaudalis | 0.66 | 0.19 | 0.99 | 0.81 | 0.63 | 0.18 |
| Tibialis Anterior | 0.66 | 0.99 | 0.87 | 0.63 | 0.24 | 0.90 |

<sup>\*</sup>*P*-values were derived from two-way analysis of variance followed by Tukey's multiple comparisons test, with the significance levels set to 5%.

We then measured the length of the muscle fibers (L_f_). Even though L_s_ did not differ between the groups, we additionally controlled for any potential differences between specimens at the time of fixation. To this effect, we calculated normalized fiber length (L_fn_) that takes into account L_s_ within each specimen at the time of fixation, using previously established methods (L_fn_ = S_n_ x L_so,_ where S_n_ is sarcomere number (S_n_ = L_f_/L_s_) and L_so_ is species-specific optimal L_s_ (2.4 µm in rat).^18^ For the reduced load/pregnancy hormones^+^ group, we compared PFM L_fn_ between the sides with and without conceptuses. The differences between the contralateral sides were observed in coccygeus and pubocaudalis, with L_fn_ on the side ipsilateral to the uterine horn with conceptuses significantly exceeding that on the side with ligated uterine horn in coccygeus (*P* = 0.03) and approaching statistical significance in pubocaudalis (*P* = 0.07, Figure 3). There were no differences between contralateral sides for iliocaudalis or tibialis anterior, *P* > 0.9. We, therefore, used the values from the side ipsilateral to the ligated uterine horn for comparisons across the experimental groups.

**Figure 3.**
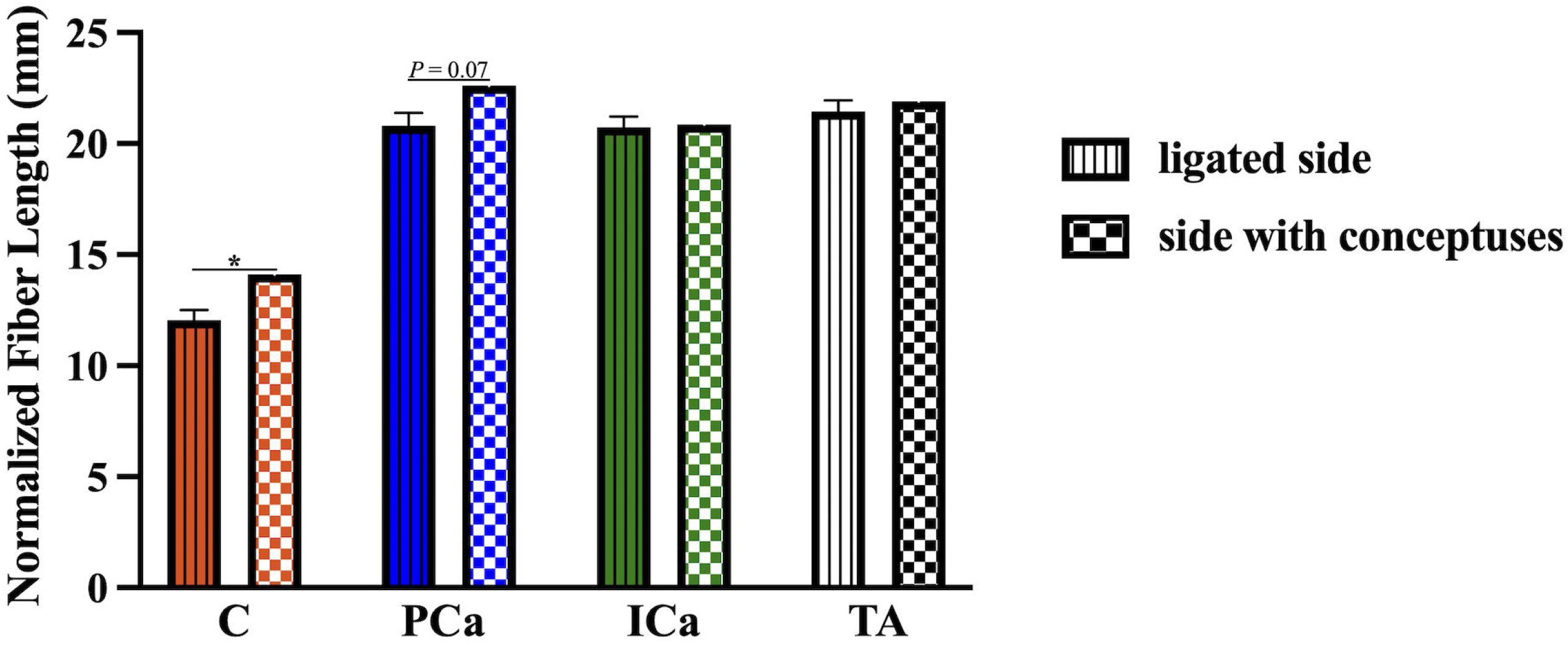
Comparison of pelvic floor muscle normalized fiber lengths (in millimeters) between side with conceptuses and side without conceptuses (ligated uterine horn) in reduced load/hormones^+^ group. *Footnote:* **C**, coccygeus; **PCA**, pubocaudalis; **ICa**, iliocaudalis; **TA**, tibialis anterior. Data are presented as mean ± standard error of mean. Some standard errors of mean values were too small to be visible as error bars. \**P* < 0.05 derived from paired Student’s t-test.

The following results are presented in comparison to the load^-^/pregnancy hormones^-^ control group, unless stated otherwise (Table 2). Coccygeus demonstrated addition of sarcomeres in series in response to muscle and sarcomere stretch induced by the increased load uncoupled from pregnancy hormones, as well as reduced load in the presence of pregnancy hormones. The above is evident from the increased L_fn_ in both the load^+^/pregnancy hormones^-^ (*P*<0.0001) and reduced load/pregnancy hormones^+^ groups (*P*=0.01). Moreover, coccygeus L_fn_ in these groups did not differ from that observed in the load^+^/pregnancy hormones^+^ group (*P*>0.5). In contrast, pubocaudalis L_fn_ increased significantly in the load^+^/pregnancy hormones^-^ (*P*<0.05), but not in the reduced load/pregnancy hormones^+^ (*P*>0.1). For iliocaudalis, substantial sarcomerogenesis occurred in response to non-reduced load and pregnancy hormones together, as evident by significant increase in L_fn_ only in the load^+^/pregnancy hormones^+^ group (*P*<0.05). In contrast to PFMs, tibialis anterior was not affected by either the increased load, pregnancy hormones, or the combination of these physiological cues (*P*>0.1).

**Table 2.** Normalized fiber length (L_fn_, millimeters) of the pelvic floor muscles and tibialis anterior in the presence of load and/or pregnancy hormones presented as mean ± standard error of mean

| Muscle | Load<sup>-</sup>/<br>Hormones<sup>-</sup><br>(N=10) | Load<sup>+</sup>/<br>Hormones<sup>-</sup><br>(N=9) | *P*-<br>value<sup>*</sup> | Reduced<br>Load/<br>Hormones<sup>-</sup><br>(N=9) | *P*-<br>value<sup>*</sup> | Load<sup>+</sup>/<br>Hormones<sup>+</sup><br>(N=5) | *P*-<br>value<sup>*</sup> |
| --- | --- | --- | --- | --- | --- | --- | --- |
| Coccygeus | 9.97 ± 0.26 | 13.33 ± 0.94 | <0.0001 | 12.06 ± 0.44 | 0.0075 | 12.82 ± 0.30 | 0.0018 |
| Pubocaudalis | 19.52 ± 0.34 | 21.20 ± 0.52 | 0.0406 | 20.80 ± 0.59 | 0.1567 | 22.53 ± 0.33 | 0.0009 |
| Iliocaudalis | 19.45 ± 0.42 | 21.00 ± 0.46 | 0.0646 | 20.74 ± 0.49 | 0.1540 | 21.45 ± 0.83 | 0.0407 |
| Tibialis<br>Anterior | 20.07 ± 0.39 | 20.58 ± 0.44 | 0.8076 | 21.44 ± 0.51 | 0.1202 | 21.21 ± 0.39 | 0.3770 |
|  | *P*-value<sup>†</sup> |  |  |  |  |  |  |
| Muscle | Load<sup>-</sup>/<br>Hormones<sup>-</sup><br>vs<br>Load<sup>+</sup>/<br>Hormones<sup>+</sup> | Load<sup>-</sup>/<br>Hormones<sup>-</sup><br>vs<br>Load<sup>+</sup>/<br>Hormones<sup>-</sup> | Load<sup>-</sup>/<br>Hormones<sup>-</sup><br>vs<br>Reduced<br>Load/<br>Hormones<sup>+</sup> | Load<sup>+</sup>/<br>Hormones<sup>+</sup><br>vs<br>Load<sup>+</sup>/<br>Hormones<sup>-</sup> | Load<sup>+</sup>/<br>Hormones<sup>+</sup><br>vs<br>Reduced<br>Load/<br>Hormones<sup>+</sup> | Load<sup>+</sup>/<br>Hormones<sup>-</sup><br>vs<br>Reduced<br>Load/<br>Hormones<sup>+</sup> |  |
| Coccygeus | 0.003 | <0.0001 | 0.01 | 0.93 | 0.79 | 0.27 |  |
| Pubocaudalis | 0.002 | 0.07 | 0.24 | 0.38 | 0.16 | 0.94 |  |
| Iliocaudalis | 0.07 | 0.11 | 0.24 | 0.95 | 0.82 | 0.98 |  |
| Tibialis<br>Anterior | 0.50 | 0.88 | 0.19 | 0.87 | 0.99 | 0.61 |  |

\**P*-values were derived from two-way analysis of variance followed by Dunnett's multiple comparisons test, with the significance levels set to 5%.
†*P*-values were derived from two-way analysis of variance followed by Tukey's multiple comparisons test, with the significance levels set to 5%.

Next, we compared the ECM collagen content of PFMs and tibialis anterior subjected to the same experimental conditions. The intramuscular collagen content of coccygeus and pubocaudalis was significantly greater in the load^+^/pregnancy hormones^-^ group and the reduced load/pregnancy hormones^-^ group than in the load^-^/pregnancy hormones^-^ group, (*P* < 0.05, Figure 4). Moreover, the collagen content of coccygeus and pubocaudalis in these groups did not differ from that observed in the load^+^/pregnancy hormones^+^ group (*P*>0.5). These data indicate that load or pregnancy hormones can induce the ECM remodeling in these PFMs, with no additional increase in the intramuscular collagen observed in the presence of both cues. In the iliocaudalis and tibialis anterior, there were no differences in collagen content between any of the experimental groups (*P*>0.2).

**Figure 4.**
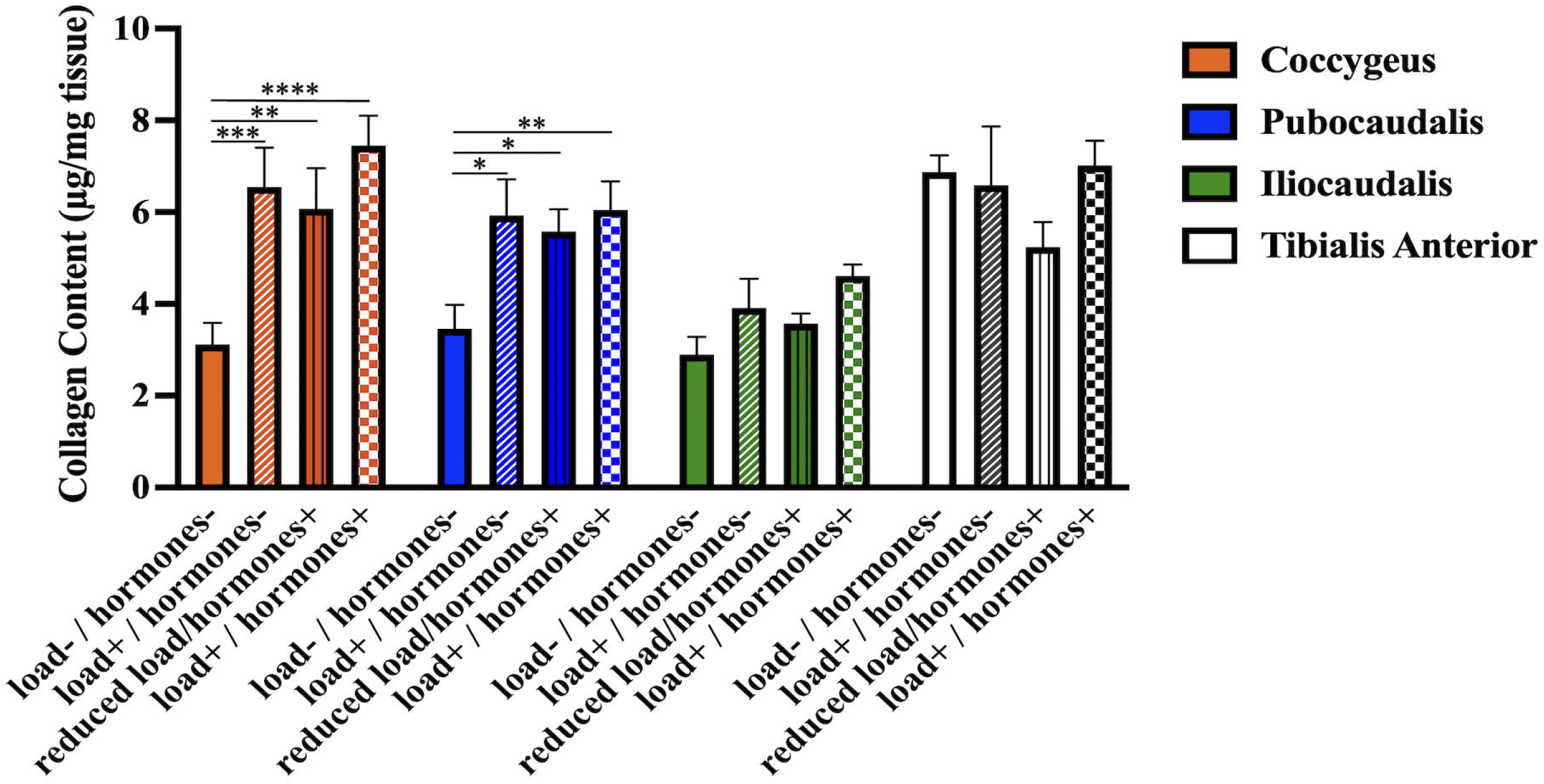
Changes in intramuscular collagen content (in micrograms per milligram of muscle tissue) in response to the independent and combinatorial effect of mechanical load and pregnancy hormone milieu as determined by measurement of intramuscular hydroxyproline content *Footnote:* Data are presented as mean ± standard error of mean. *P*-values were derived from repeated measures two-way analysis of variance followed by pairwise comparisons with Dunnett’s and Tukey’s tests. ^*^*P* < 0.05; \*\**P* < 0.01; \*\*\**P <* 0.001; \*\*\*\**P* < 0.0001

### The effect of sustained mechanical load, uncoupled from the endocrine milieu of pregnancy, on the PFMs’ response to parturition-related strains

We have previously shown that pregnancy-induced adaptations protect PFMs against birth injury, as indicated by the absence of sarcomere hyperelongation, a major cause of mechanical muscle injury, in response to parturition-related strains.^19^ To determine whether sarcomerogenesis of coccygeus and pubocaudalis induced by increased load was similarly protective against parturition-related strains, we compared the impact of vaginal distention of various magnitude between three experimental conditions (load^-^/pregnancy hormones^-^, load^+^/pregnancy hormones^-^, and load^+^/pregnancy hormones^+^). The data from historic controls were used for the load^-^/pregnancy hormones^-^ and load^+^/pregnancy hormones^+^ conditions,^19^ given confirmed reproducibility of our vaginal distention model.^25^

### Response to physiologic parturition-related strains

In response to vaginal distension with the 3mL balloon volume (physiologic strain, Figure 5A), Ls of coccygeus and pubocaudalis in the load^+^/hormones^-^ group were substantially shorter than L_s_ in the load^-^/hormones^-^ group (P<0.01). However, sarcomere elongation was still significantly longer than that in the load^+^/hormones^+^ group (P<0.001). Taken together, these data indicate that adaptations induced by increased load in the absence of hormonally driven alterations confer an intermediate protective effect against mechanical muscle injury caused by parturition-associated strains. As expected, there were no differences in iliocaudalis L_s_ between the groups (*P*>0.2), as this PFM experiences smaller strains during vaginal balloon distention.^19^

**Figure 5.**
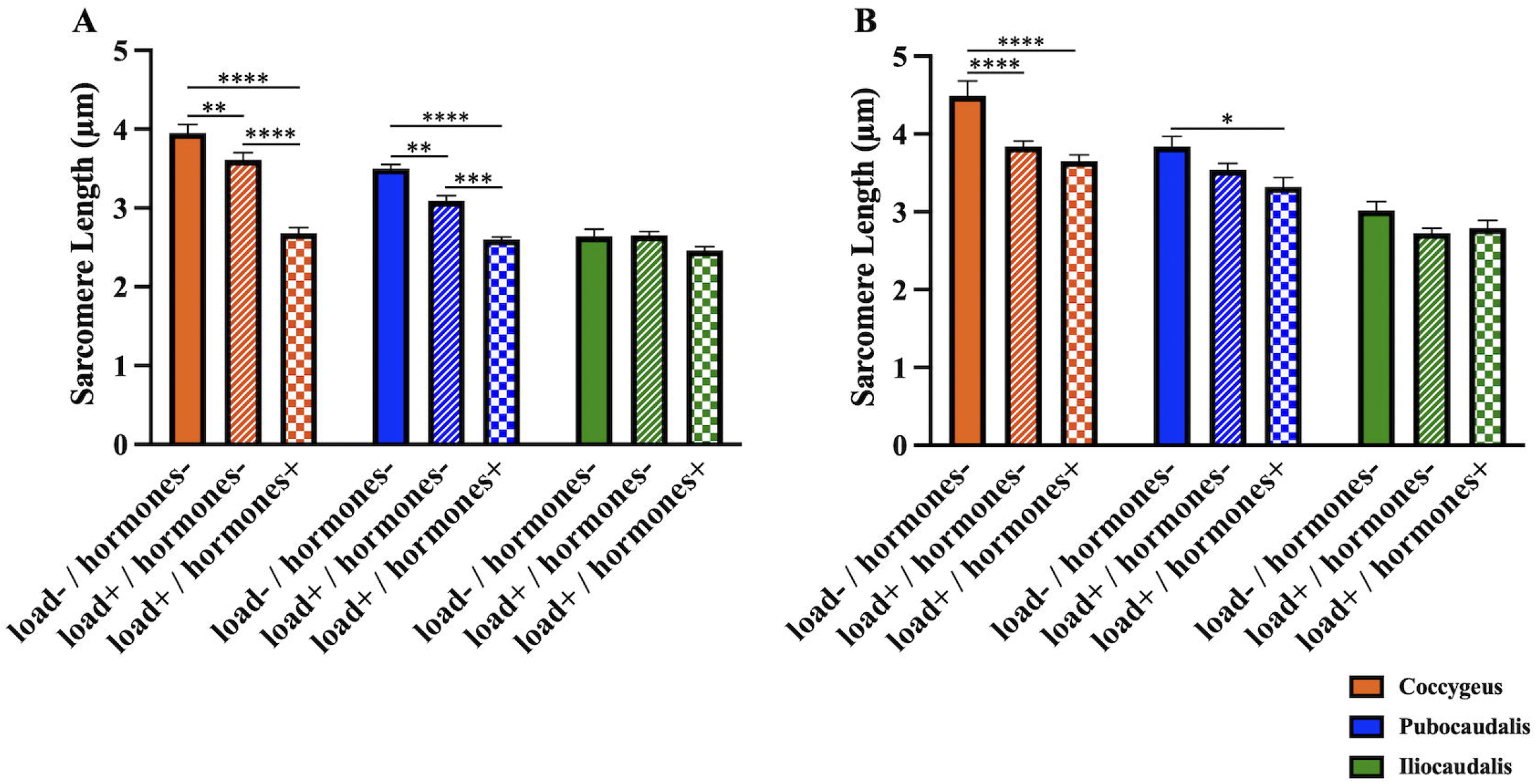
(A) Sarcomere length measurements (in micrometers) of rat pelvic floor muscles at 3 mL (physiologic) strain via vaginal distension in load^-^/hormones^-^, load^+^/hormones^-^, load^+^/hormones^+^ groups (B) Sarcomere length measurements (in micrometers) of rat pelvic floor muscles at 5 mL (supraphysiologic) strain via vaginal distension in load^-^/hormones^-^, load^+^/hormones^-^, load^+^/hormones^+^ groups Footnote: Data are presented as mean ± standard error of mean. *P*-values were derived from repeated measures two-way analysis of variance followed by pairwise comparisons with Tukey’s test. \**P* < 0.05; \*\**P* < 0.01; \*\*\**P <* 0.001; \*\*\*\**P* < 0.0001

### Response to supraphysiologic parturition-related strains

Next, we determined whether mechanical load provided protection against supraphysiologic strains (Figure 5B). Like the response to vaginal distention with 3mL volume, coccygeus L_s_ in the load^+^/hormones^-^ group was significantly shorter than the load^-^/pregnancy hormones^-^ group. However, as opposed to the smaller strain, L_s_ in the load^+^/hormones^-^ group did not differ from L_s_ in the load^+^/ hormones^+^ group (*P*>0.5). With respect to pubocaudalis, L_s_ in load^+^/hormones^-^ group did not differ from that in either load^-^/pregnancy hormones^-^ (*P*>0.1) or the load^+^/hormones^+^ (*P*>0.2) group. This confirms that our model reproduced adaptations that are protective against physiological strains, and that the protective effect of these adaptations diminishes when mechanical insult associated with parturition is excessive. Iliocaudalis L_s_ also did not differ between the groups (*P*>0.1).

## COMMENT

### Principal Findings

The plasticity of the individual components of the rat PFM complex in response to mechanical and endocrine cues is variable. Out of all PFMs, coccygeus is the most susceptible to either stimuli, with sarcomerogenesis observed in all experimental conditions tested (load^+^/pregnancy hormones^-^, reduced load/pregnancy hormones^+^, load^+^/pregnancy hormones^+^) compared to the unperturbed controls (load^-^/pregnancy hormones^-^). In pubocaudalis, fiber length increased in response to load alone and in combination with pregnancy hormones. Fiber length of iliocaudalis increased only in response to the combinatorial effect of mechanical and hormonal cues. These results indicate that coccygeus responds to either mechanical or endocrine stimulus. With respect to pubocaudalis, loading is sufficient to induce sarcomerogenesis and mechanical cue is likely the dominant driver of this adaptation in this portion of the rat levator ani muscle. The key role of mechanical load in the plasticity of the contractile component of these muscles is further supported by our findings in the unilaterally pregnant (reduced load/hormones^+^) group. L_fn_ on the side ipsilateral to the uterine horn with conceptuses was significantly greater than on the side with ligated uterine horn in coccygeus and pubocaudalis, where this difference approached statistical significance. Importantly, the extent of PFM elongation by sarcomerogenesis in coccygeus and pubocaudalis in the load^+^/hormones^-^ group was equivalent to that observed in the unperturbed pregnant (load^+^/hormones^+^) rats. On the other hand, neither load alone nor hormonal stimulation with reduced load are sufficient to induce sarcomerogenesis of iliocaudalis, which required both mechanical and endocrine cues. In contrast to PFMs and consistent with our previous findings,^18^ the hind limb tibialis anterior muscle was not affected by either the increased mechanical load imposed by the weighted uterine horns, pregnancy hormones, or the combination of these physiological cues, suggesting that PFMs are uniquely and differentially susceptible to these perturbations.

Like the response of the contractile myofibers, intramuscular ECM remodeling induced by mechanical load in the presence or absence of pregnant hormonal milieu varied across individual PFMs. Intramuscular collagen content of coccygeus and pubocaudalis increased in response to either load or hormonal stimuli. As with PFM elongation by sarcomerogenesis, the increase in ECM collagen content of coccygeus and pubocaudalis in the load^+^/hormones^-^ group was equivalent to that observed in the pregnant rats (load^+^/hormones^+^). We did not observe an increase in collagen content in iliocaudalis or tibialis anterior in any of the experimental conditions compared to the load^-^/hormones^-^ controls. These results indicate that, as with sarcomerogenesis, ECM remodeling of coccygeus and pubocaudalis is induced by either mechanical or endocrine stimulus.

The importance of pregnancy-induced adaptations in the pelvic soft tissues mainly lies in their protective function against maternal birth injury. To this effect, we examined the response of chronically loaded PFMs to parturition-associated strains of various magnitudes. We found that adaptations of coccygeus and pubocaudalis, resultant from increased load, conferred protective effect against mechanical muscle injury relative to the response of the control muscles (load^-^/hormones^-^). However, this protective effect was smaller than that afforded by the adaptative changes of PFMs exposed to load and pregnant hormonal milieu. With respect to the supraphysiological strains, adaptations of coccygeus induced by load alone are sufficient to protect against mechanical muscle injury, based on our finding that sarcomere length in the load^+^/hormones^-^ group was significantly less than the hyperelongated sarcomeres in the unperturbed load^-^/hormones^-^ group and not different from the load^+^/hormones^+^ group. Load-induced adaptations of pubocaudalis were inadequate to confer protection against sarcomere hyperelongation, when this muscle experienced a higher magnitude strains.

### Results in the Context of What is Known

Taken together, our results support the hypothesis that initial sarcomere elongation promotes sarcomerogenesis of PFMs, ultimately leading to return of sarcomeres to their operational length, which is necessary for optimal *in vivo* muscle function. Prior investigations in various animal models have demonstrated a similar phenomenon in limb muscles – that muscles placed under chronic stretch elongate via sarcomerogenesis.^26,27^ Overall, increased mechanical load appears to play a key role in driving pregnancy-induced adaptations in the rat PFMs. In our study, load-induced sarcomerogenesis and increase in the intramuscular collagen content in the non-pregnant model varied by muscle, with coccygeus most responsive (alterations with either load or pregnancy hormones), iliocaudalis least responsive (alterations with neither load nor pregnancy hormones alone), and pubocaudalis demonstrating responsiveness intermediate to coccygeus and iliocaudalis. Importantly, PFM plasticity in response to increased load afforded protection against PFM mechanical birth injury, with degree of protection varying between PFMs and strain magnitude. Taken together, this points towards a differential sensitivity of the individual pelvic skeletal muscles to the physiological cues associated with pregnancy. In addition, our findings suggest that the combinatorial effect of the endocrine and mechanical signals plays an important role in the PFM response to parturition-related strains.

### Clinical Implications

Our findings suggest that modulation of the PFM stretch induced by mechanical load during pregnancy, such as with specific pelvic floor training regimens, may be a potential therapeutic intervention for augmenting the protective antepartum PFM plasticity and preventing muscle injury during vaginal delivery.

### Research Implications

In the current study we begin to elucidate the multifactorial mechanisms that govern PFM plasticity during pregnancy. Determining the key drivers of the protective adaptations of PFMs in the pre-clinical model is essential for promoting our understanding of the potential PFM plasticity in pregnant women and its role in modulating one’s predisposition to birth injury. To date, the causes underlying differential pelvic soft tissue damage during parturition and subsequent development of pelvic floor disorders in vaginally parous women remains unknown, and no effective strategies exist for the prevention of maternal birth injury.

### Strengths and Limitations

The strengths of the current work include the use of the rat model, specifically validated for the studies of the human PFMs^13,14^; the development of the novel non-pregnant rat model of PFM mechanical loading; and the first evaluation of the role of mechanical load and related muscle stretch, in the presence and absence of the endocrine cues of pregnancy, in PFM plasticity.

The limitations of our study are inherent to the use of experimental models to simulate human condition. However, direct PFM tissue studies are not possible in asymptomatic living women, thus precise tissue-level experiments in the animal model are the necessary step in the continuum of these clinically relevant studies. Also, given a reduction in placentally-derived factors in rats with a smaller number of gestations, the hormonal milieu in the reduced load/pregnancy hormones^+^ may be different compared to animals with a larger number of conceptuses. Unfortunately, currently there is no known way to induce the complete spectrum of endocrine changes that occur in pregnancy without the pregnancy itself, which constitutes a load on PFMs. Also, we were unable to steadily increase the mechanical load over time as occurs in a normal gestation; instead, PFMs were exposed to the same load for the entire 21 days. Therefore, the effects of mechanical load in pregnancy could be overestimated. However, the extent of sarcomerogenesis and increase in PFM ECM collagen content were comparable in our non-pregnant loaded model and pregnant animals. Finally, since we based our *a priori* sample size calculation on differences between the non-pregnant (load^-^/hormones^-^) and pregnant (load^+^/hormones^+^) controls, we performed *post hoc* power analysis and found that we only had 48% power to detect a difference in PCa L_fn_ between reduced load/hormones^+^ and non-pregnant controls.

## Conclusions

Load induces plasticity of the intrinsic pelvic floor muscle components that renders protection against mechanical birth injury. The protective effect, which varies between individual muscles and strain magnitudes, is further augmented by the presence of pregnancy hormones. Maximizing impact of mechanical load on pelvic floor muscles during pregnancy, such as with specialized pelvic floor muscle stretching regimens, is a potentially actionable target for augmenting pregnancy-induced adaptations to decrease birth injury in women who may otherwise have incomplete antepartum muscle adaptations.

## ACKNOWLEDGEMENTS

We gratefully acknowledge our funding source – NIH grant R01 HD092515 from Eunice Kennedy Shriver National Institute of Child Health and Human Development-that supported this project.

---

**Supplemental Table 1.** Normalized fiber length (L_fn_, millimeters) of the pelvic floor muscles and tibialis anterior on loaded and non-loaded side in the Load^+^/Hormones^-^ group presented as mean ± standard error of mean

| Muscle | Non-loaded side (N=9) | Loaded side (N=9) | *P*-value* |
| --- | --- | --- | --- |
| Coccygeus | 12.88±0.54 | 13.33±0.94 | 0.9763 |
| Pubocaudalis | 21.57±0.62 | 21.19±0.52 | 0.8743 |
| Iliocaudalis | 20.22±0.75 | 21.00±0.46 | 0.8478 |
| Tibialis Anterior | 21.79±0.41 | 20.58±0.44 | 0.5197 |

\**P*-values were derived from two-way analysis of variance followed by Šídák's multiple comparisons test, with the significance levels set to 5%.

